# Rare, pathogenic germline variants in *Fanconi Anemia* genes increase risk for squamous lung cancer

**DOI:** 10.1101/269837

**Authors:** Myvizhi Esai Selvan, Robert J. Klein, Zeynep H. Gümüş

**Affiliations:** Department of Genetics and Genomic Sciences; Icahn Institute for Genomics and Multiscale Biology, Icahn School of Medicine at Mount Sinai, New York, New York, USA.

## Abstract

**Purpose:** Lung cancer is the leading cause of cancer deaths worldwide, with substantially better prognosis in early stage as opposed to late stage disease. Identifying genetic factors for lung squamous carcinoma (SqCC) risk will enable their use in risk stratification, and personalized intensive surveillance, early detection, and prevention strategies for high-risk individuals.

**Study Design and Participants:** We analyzed whole-exome sequencing datasets of 318 cases and 814 controls (discovery cohort) and then validated our findings in an independent cohort of 444 patients and 3,479 controls (validation cohort), all of European descent, totaling a combined cohort of 765 cases and 4,344 controls. We focused on rare pathogenic variants found in the ClinVar database and used penalized logistic regression to identify genes in which such variants are enriched in cases. All statistical tests were two-sided.

**Results:** We observed an overall enrichment of rare, deleterious germline variants in *Fanconi Anemia* genes in cases with SqCC (joint analysis OR=3.08, *p*=1.4e-09, 95% confidence interval [CI]=2.2–4.3). Consistent with previous studies, *BRCA2* in particular exhibited an increased overall burden of rare, deleterious variants (joint OR=3.2, *p*=8.7e-08, 95% CI=2.1–4.7). More importantly, rare deleterious germline variants were enriched in *Fanconi Anemia* genes even without the *BRCA2* rs11571833 variant that is strongly enriched in lung SqCC cases (joint OR=2.76, *p*=7.0e-04, 95% CI=1.6–4.7).

**Conclusions:** These findings can be used towards the development of a genetic diagnostic test in the clinic to identify SqCC high-risk individuals, who can benefit from personalized programs, improving prognosis.

## Introduction

Lung cancer is the leading cause of cancer mortality in the United States (1). Although the disease is significantly more common in individuals with a positive family history (Odds Ratio (OR) 1.57-5.52) (2,3) and has an estimated heritability of 18% (4), genome-wide scans have only identified common loci associated with modest risk (5). Identifying individuals at increased genetic risk for lung cancer would lead to personalized surveillance programs for diagnosis and improve prognosis. Recent studies have identified rare variants in *BRCA2* and *CHEK2* genes with large effects in genome-wide scans of lung cancer patients (6). Interestingly, seemingly sporadic cancer patients also carry a significant number of rare germline variants of unknown significance (7,8).

Here, we utilized whole exome sequencing (WES) data on a lung cancer case-control cohort from Transdisciplinary Research in Cancer of the Lung (TRICL) (http://u19tricl.org) to examine the role of rare coding variants in risk for lung squamous cell carcinoma (SqCC), and validated our results in an independent case-control cohort by utilizing the untapped resources of cases in The Cancer Genome Atlas (TCGA) (http://cancergenome.nih.gov) and controls from various studies in database of Genotypes and Phenotypes (dbGaP) (http://www.ncbi.nlm.nih.gov/gap). We confirm previous observations regarding the role of *BRCA2* in lung cancer risk and expand this observation and identify an overall burden of rare, deleterious variants in *Fanconi Anemia (FA)* genes, and especially those in the *FA* core complex, in lung SqCC. While *FA* genes have been previously implicated in cancer risk (9), ours is the first report of a role for them in lung cancer risk.

## Methods

### Data Sources

We downloaded data for two case-control cohorts. The discovery case-control cohort was from the Transdisciplinary Research in Cancer of the Lung (TRICL) project, which we downloaded from dbGaP (phs000876). The validation cases were from The Cancer Genome Atlas (TCGA) and controls were from eight population-based studies in dbGaP. We downloaded TCGA germline whole exome sequencing bam files from National Cancer Institute Genomic Data Commons (GDC) data portal (https://gdc-portal.nci.nih.gov). We downloaded the control samples from dbGaP studies: Multiethnic Study of Atherosclerosis (MESA) cohort (phs000209), STAMPEED study: Northern Finland Birth Cohort (NFBC) 1966 (phs000276), NHLBI GO-ESP: Lung Cohorts Exome Sequencing Project (COPDGene) (phs000296), Common Fund (CF) Genotype-Tissue Expression (GTEx) (phs000424), Genetic Analyses in Epileptic Encephalopathies: A sub-study of Epi4K - Gene Discovery in 4,000 Epilepsy Genomes: (phs000654), ARRA Autism Sequencing Collaboration (phs000298), Bulgarian schizophrenia trio sequencing study (phs000687) and Myocardial Infarction Genetics Exome Sequencing (MIGen_ExS) Consortium: Ottawa heart study (phs000806).

### Study cohorts

We first analyzed the discovery cohort of 333 familial-enriched lung SqCC cases and 853 controls, then the validation cohort of 494 sporadic cases and 3,954 controls. The combined cohort included 827 cases and 4807 controls. All cohorts and their sample sizes are listed in Supplementary Table S1.

### Variant discovery

We performed variant discovery by realignment and joint analysis of all case and control germlines using GVCF-based best practices for the Genome Analysis Toolkit (GATK, https://www.broadinstitute.org/gatk/) as implemented in a custom pipeline at the Icahn School of Medicine at Mt. Sinai (10).

### Sample QC

We removed samples with <85% missing data. We identified duplicates and related individuals by first or second degree based on analysis with KING software (11), where we removed a sample from each inferred pair that had a higher fraction of missing data. We removed any bias that may arise due to systematic ancestry-based variations in allele frequency differences between cases and controls by using samples with European ancestry as described below.

### Population stratification

To remove spurious associations and to adjust for population stratification, we used Principle Component Analysis (PCA). To identify the population structure, we removed indels and rare variants (defined by <5% of minor allele frequency, MAF), using 1000 Genomes dataset (12) and The Ashkenazi Genome Consortium (TAGC) as reference (https://ashkenazigenome.org). For the remaining variants, we performed Linkage Disequilibrium (LD) pruning, filtered for a call rate of at least 0.99, and did PCA with smartpca in EIGENSOFT 5.0.1 software. We filtered for the least ancestry-based variation by focusing our downstream analyses on the largest set of individuals clustered within the PCA plot by PCA gating. As shown in PCA plots of the three case-control cohorts along with the gated regions in Fig. 1, these correspond to European ancestry.

**Figure 1:**
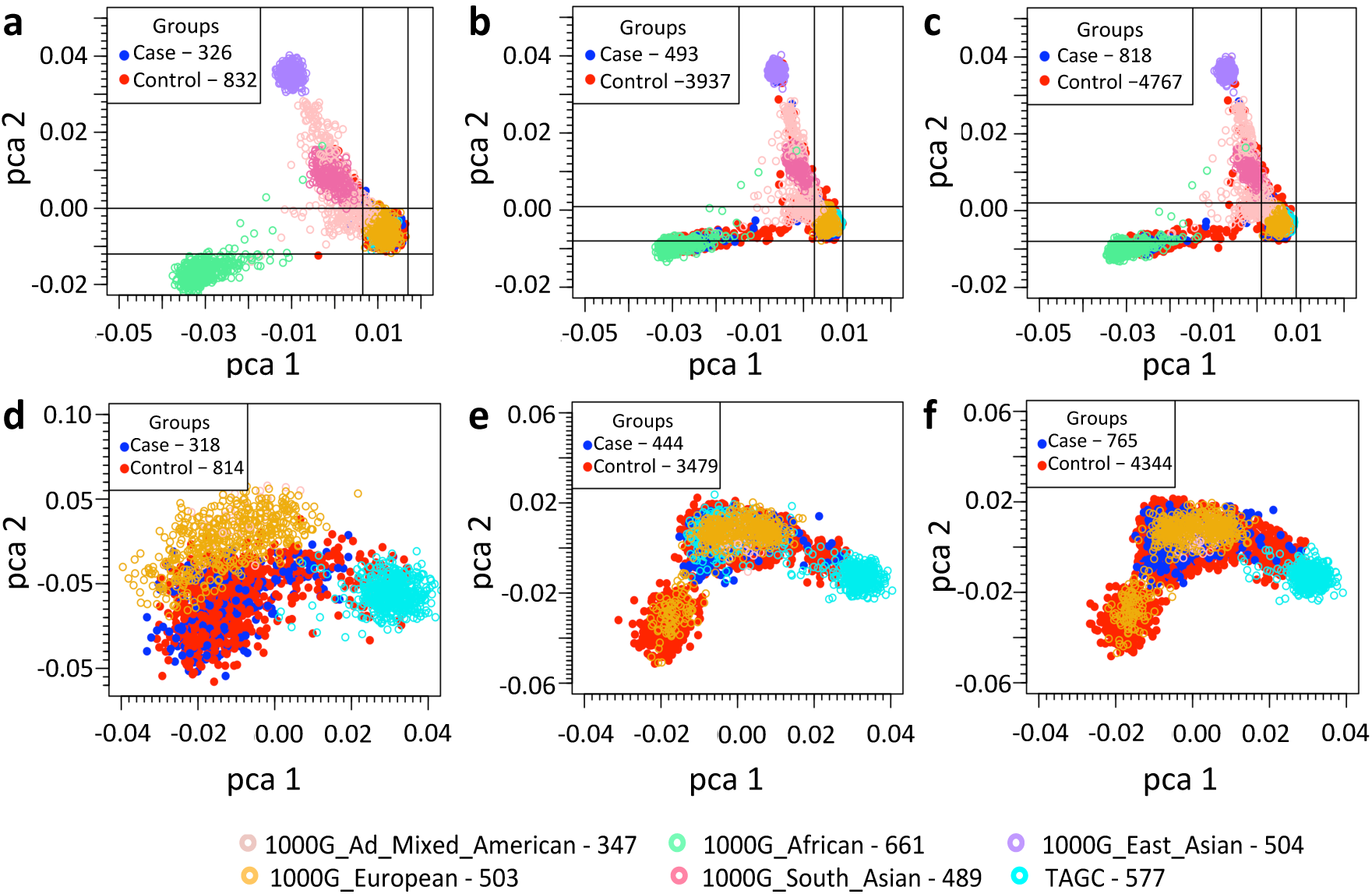
Principal Component Analyses (PCA) of all study cohorts and all gated study cohorts. PCA based on common SNPs (MAF > 0.05) showing the top two principal components of (i) the study cohorts together with 1000 Genomes and TAGC samples (a-c) and of (ii) the gated samples from the study cohorts with European ancestry (d-f). **a)** Discovery cohort (TRICL); **b)** Validation cohort (TCGA-dbGaP); **c)** Combined cohort (TRICL-TCGA-dbGaP); **d)** Gated samples of discovery cohort (318 cases and 814 controls)**; e)** Gated samples of validation cohort (444 cases and 3479 controls); **f)** Gated samples of combined cohort (765 cases and 4344 controls).

### Variant-level QC

Within the individuals that passed sample QC, we focused on ensuring high-quality genotype/variant calls for analysis. For this purpose, we filtered for variants with: read genotype quality ≥20; read depth ≥10; allelic depth of alternate allele ≥4; variant sites with quality score ≥50; quality by depth score ≥2; mapping quality ≥40; read position rank sum >–3; mapping quality rank sum >–10 and variant tranche <99%. For heterozygous genotypes, we filtered for alternative allele ratio between 0.30 and 0.70. Finally, we kept sites with ≥ 88% of data in both cases and controls.

### Variant filtering

Among the variants that passed QC, we focused on rare variants with known pathogenic effects. Such causal variants have been shown to significantly alter encoded protein’s function and to occur significantly less frequently in control populations. To filter out common polymorphisms, we removed variants present in both cases and controls at MAF >2% or in Exome Aggregation Consortium (ExAC) non-TCGA Non-Finnish European population at MAF >1%. We considered variants that passed these filters as rare. We then filtered the rare variants for functional impact based on those present in the ClinVar database (13) using Annovar (http://annovar.openbioinformatics.org) (versions: 2016Feb01, clinvar_20170905). We considered a variant as pathogenic if it was: (i) listed pathogenic/likely pathogenic in ClinVar; or (ii) a frameshift or stopgain variant located 5’ of a listed pathogenic LOF variant (nonsense or frameshift) in ClinVar. We did not consider missense variants with conflicting pathogenicity in ClinVar (13) (where some submissions state pathogenic/likely pathogenic and others state benign/likely benign/uncertain significance). To reduce case-control sample differences, we kept sites with differential missing variant fraction ≤ 0.05 between the cases and controls.

### Gene-sets

We considered six gene-sets associated with DNA repair and cancer predisposition (CPD) for gene-set level burden analysis of rare, pathogenic variants associated with lung SqCC risk, with full details in Supplementary Table S2.

### Statistical analysis

#### Background variation correction

To test for possible background variation, we calculated the tally of rare autosomal synonymous variants per individual between cases and controls. We defined synonymous variants as rare at Exac MAF ≤0.005% and MAF ≤0.05% in combined case-control cohort. Supplementary Fig. S1 provides the distribution and background variation statistics of genes with rare synonymous variants in all cohorts. We accounted for differences in background variation as described below.

#### Variant burden analyses

We identified risk genes associated with lung SqCC based on aggregate rare, pathogenic variant burden for each gene using Penalized Logistic Regression Analysis (PLRA), using the logistf package in R (https://cran.r-project.org/web/packages/logistf/index.html). To adjust for background variation, we used the number of genes with rare synonymous variants as a covariate for each individual. We filtered out genes with minimal number of pathogenic variants (cases ≤2 and controls ≤1). We deemed genes with *p*-value ≤0.05 statistically significant. For gene-set level burden analysis, instead of considering the number of rare pathogenic variants in each individual, we considered if an individual has a pathogenic variant in a gene only when using PLRA. All statistical tests were two-sided.

#### Smoking effects

To account for the effects of smoking on gene burden in discovery (cases: 4.1% never smoker (NS); 95.9% smoker; controls: 34.8% NS and 65.2% smoker) and validation cohorts (cases: 2.7% NS; 95% smoker), we used PLRA with pack-years smoked as second covariate with background variation as the first covariate. For smokers with missing pack-year values in discovery cohort (1.9% cases; 1.1% controls), we imputed from average number of pack-years of other smokers. Resulting ORs and p-values of all gene-sets are in supplementary Fig. S2.

#### Age and gender effects

For the effects of age and gender on gene burden, we used PLRA with the second covariate as i) gender; ii) age; and iii) both age and gender, with background variation as the first covariate. Male and female percentages in all cohorts are provided in supplementary Table S3. We removed few control samples from analysis that had missing gender data (2 in validation and 3 in combined cohorts). Resulting ORs and p-values of all genesets are in supplementary Fig. S3-4.

## Results

### Identification of rare deleterious variants

To identify rare deleterious germline variants in individuals with lung SqCC, we utilized WES datasets from several previously published studies whose data are available through dbGaP. As the discovery cohort, we used 333 lung SqCC exomes enriched in familial cases and 853 control exomes from the TRICL study. As a validation cohort, we used 494 sporadic (non-familial) lung SqCC exomes from TCGA together with 3,954 control exomes from dbGaP. For all samples, we harmonized the data by realigning and jointly calling germline genetic variants using GATK Best Practices. We then performed QC and removed related individuals up to second degree. We also removed ancestry-based bias by classifying the genetic ancestry of all samples based on PCA and only filtering out individuals with European ancestry (Fig. 1). After sample QC, the discovery dataset included 318 cases and 814 controls and the validation included 444 cases and 3,479 controls (Supplementary Table S1). To avoid potential confounding, we asked if the frequency of neutral variation varied among cohorts (Supplementary Fig. S1) and then adjusted for differences we observed in variation (see Methods).

### Single gene analysis

We first asked if any gene showed evidence for an enrichment of rare, deleterious variants in cases compared to controls. While numerous computational approaches exist for predicting the functional impact of a missense mutation in protein-coding sequence (14), these predictions are not ready for use in the clinic. Therefore, we focused on those genes present in the ClinVar database (13), with strict pathogenicity criteria as described in Methods. When we applied these criteria to rare variants, we were able to test the remaining 593 genes after filtering in the discovery cohort.

We note that while a *BRCA2* stopgain variant, rs11571833 (p.Lys3326Ter), was previously reported to confer risk for lung SqCC (6), it did not satisfy our pathogenicity criteria as there were inconsistent reports of its pathogenicity in ClinVar (13) (29 studies report benign/likely benign; one study reports pathogenic), which annotated its clinical significance as benign. Testing this variant, we observed that results support the previous findings on its association with lung SqCC (6) risk, with a statistically significant difference between cases and controls (OR=3.0, *p*=5.3e-03, 95% CI=1.4–6.4 in discovery cohort and OR=3.3, *p*=4.4e-05, 95% CI=1.9–5.5 in validation cohort). Furthermore, among 20 individuals with this variant (12 males, 8 females) in the validation cohort, we observed Loss of Heterozygosity (LOH) in 8 (5 males, 3 females). This warrants more in-depth functional studies of this variant.

Similarly, a *CHEK2* variant, rs17879961 (p.Ile157Thr), was previously associated with reduced SqCC risk (6) but had conflicting pathogenicity reports in ClinVar (11 studies cite pathogenic/ likely pathogenic; 4 studies cite uncertain significance). Testing this variant, we did not observe a statistically significant difference between cases and controls (OR=0.30, *p*=0.11, 95% CI=0.03–1.26 in discovery cohort and OR=0.52, *p*=0.27, 95% CI=0.11–1.55 in validation cohort). In our gene-level tests, we therefore included *BRCA2* rs11571833 but not *CHEK2* rs17879961.

Testing genes with rare, pathogenic variants at gene-level, *BRCA2* was a top significant gene associated with lung SqCC in our discovery cohort (*p*=1.5e-03; OR=3.2; 95% CI=1.6–6.4), which we validated in the validation (*p=*8.0e-05; OR=3.0; 95% CI=1.8– 5.0) and combined cohorts (*p*=8.7e-08; OR=3.2; 95% CI=2.1–4.7). Without the rs11571833 variant, we still observed a statistically significant association in the combined cohort (*p*=0.04; OR=3.3; 95% CI=1.1–9.3).

Previous studies have additionally identified pathogenic germline variants in DNA damage response genes *BRCA1 (FANCS), ATM, BRIP1 (FANCJ), PALB2 (FANCN)* and *PMS2* in solid tumors in TCGA datasets (15,16) While we did not observe a statistically significant gene-level burden in cases versus controls in these genes in our cohorts, we did observe an increased frequency of rare, pathogenic variants in cases compared to controls for *PALB2* (OR=2.6, *p*=0.2, 95% CI=0.5–11.0) and *BRIP1* (OR=3.4, *p*=0.3, 95% CI=0.3–25.7) in the combined cohort. We noted that these, and *BRCA2* are *FA* genes.

### Cancer predisposition gene-set analysis

We next studied whether clinically significant variants in CPD gene-sets were correlated with increased risk of lung SqCC. We first tested 3 gene-sets:

i. 94 genes associated with CPD and currently used for clinical genetic testing in the TruSight Cancer Gene panel (17), which are predominantly tumor suppressors involved in DNA repair and cell cycle regulation;
ii. 114 CPD genes reported by Rahman (18) to have rare variants that confer high or moderate cancer risk (relative risk >2-fold) in multiple studies;
iii. 20 core DNA repair genes (DRGs) (19) associated with autosomal dominant CPD syndromes (20).

To identify additional potential CPD gene-sets, we performed a data-driven exome-wide gene-set analysis. Briefly, we tested all genes with OR>1 in our cohorts that were also in gene-sets in Molecular Signatures Database (MSigDB) (software.broadinstitute.org). In total, we tested 16,928 gene-sets for overlaps. We observed the highest overlap with the Gene Ontology *FA* nuclear complex in all cohorts (13 genes, 38.5% overlap in combined cohort). Therefore, we additionally tested (iv) all known 22 *FA*/ *FA*-like genes (21,22); (v) 9 genes in *FA* core complex; (vi) 11 *FA* genes involved in DNA repair. We list all genes for each gene-set at Supplementary Table S2.

Testing the gene-sets (i-vi), we observed higher frequency of rare, pathogenic germline variants in cases *vs.* controls for all gene-sets and statistically significant variant burden in the combined cohort (Table 1 and Fig. 2a). However, the variant burden was statistically significant in cases *vs.* controls in all cohorts only for 22 *FA* genes and 11 *FA* DNA repair genes (Fig. 2a).

**Figure 2:**
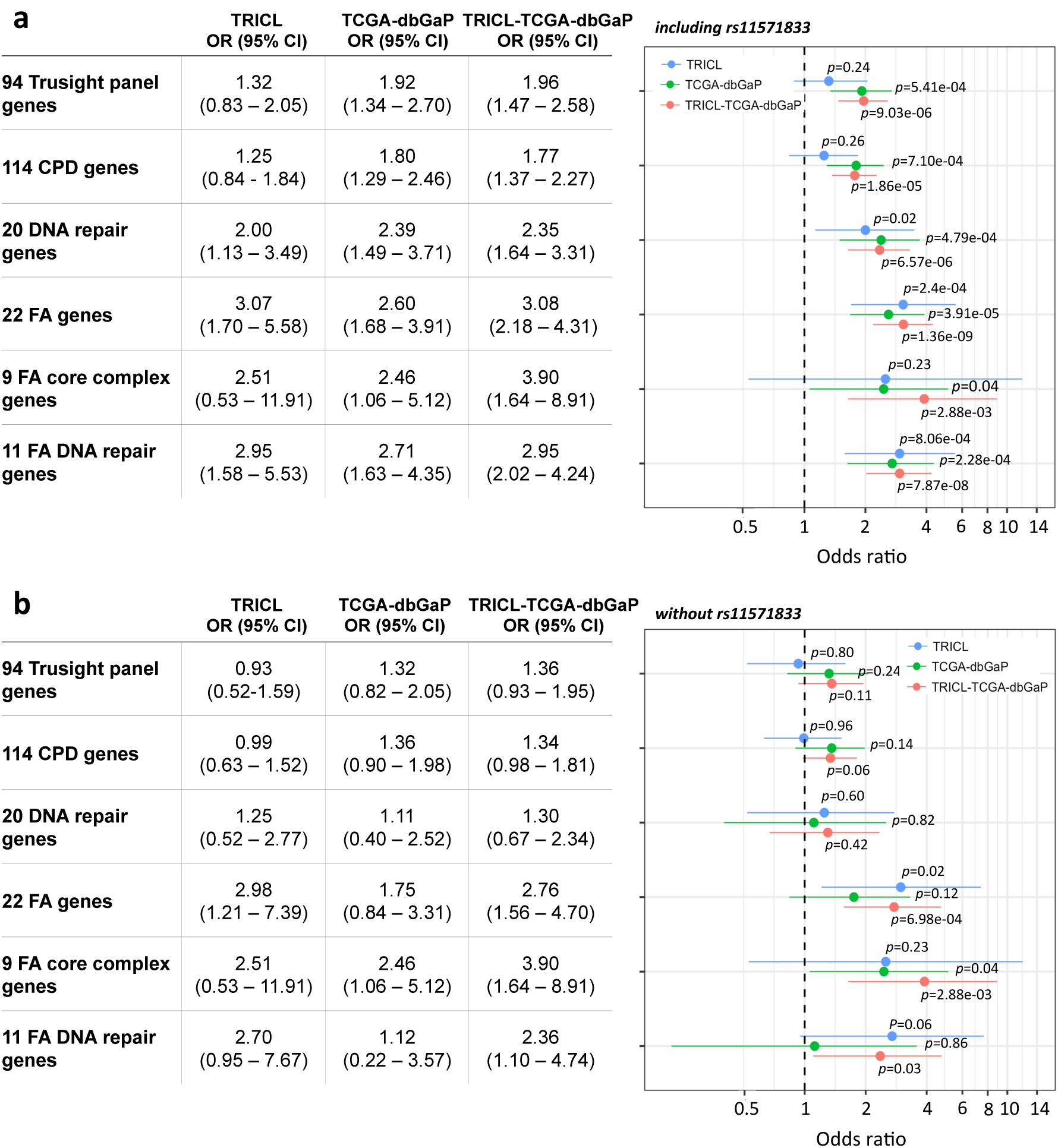
Gene-set level burden of rare, pathogenic variant p-values and Odds Ratios (ORs). For each gene-set, burden of rare, pathogenic variants for the three cohorts i) Discovery cohort (TRICL) – blue ii) Validation cohort (TCGA-dbGaP) – green iii) Combined cohort (TRICL-TCGA-dbGaP) – red **a)** including all variants; **b)** without *BRCA2* rs11571833 stopgain variant. The whiskers span the 95% confidence interval for OR values.

**Table 1:**
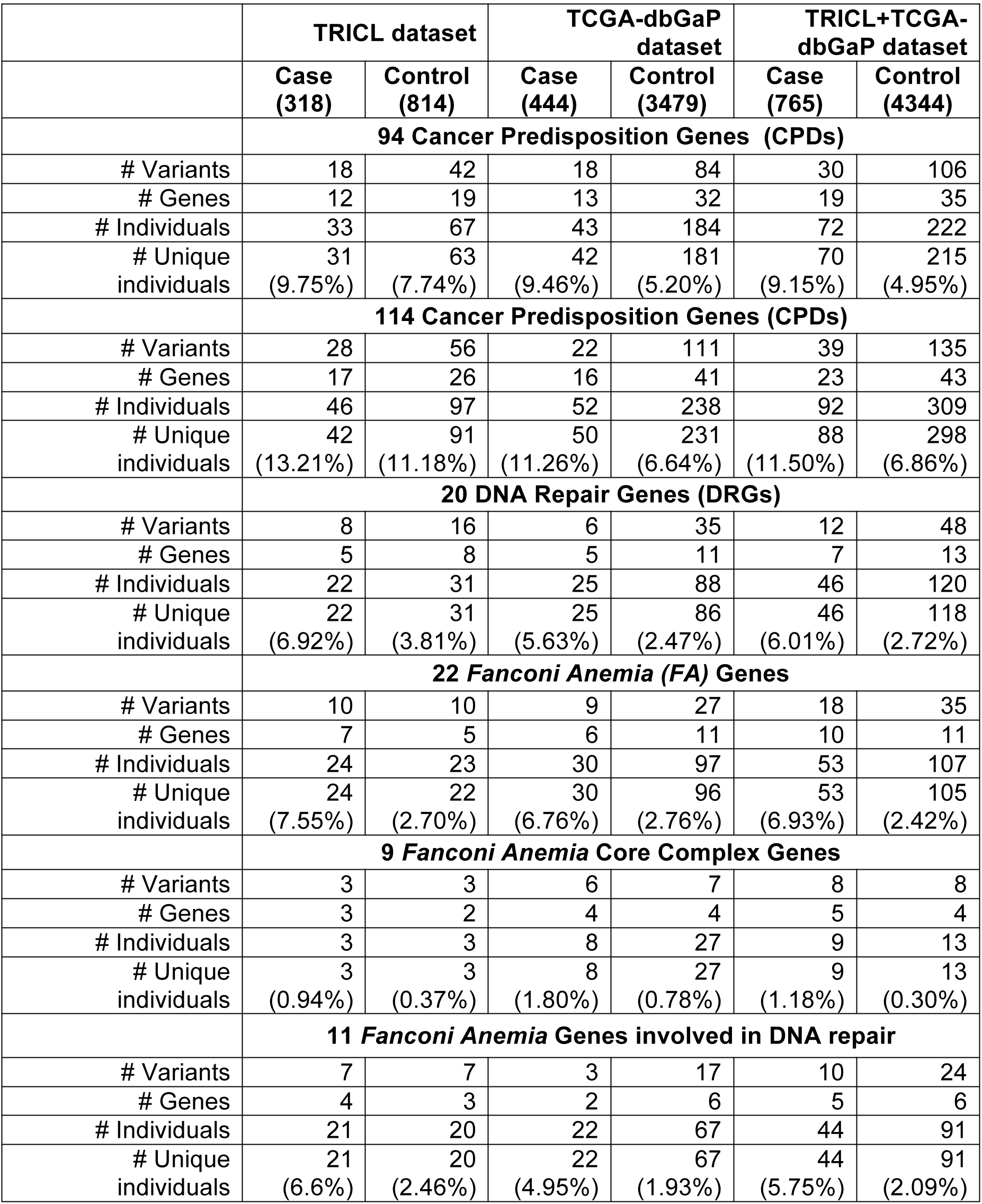
Rare, pathogenic variants in all study cohorts identified with ClinVar analysis.

To test the extent to which our findings could be influenced by the *BRCA2* rs11571833 variant, we removed it and repeated our analysis. Without this variant, we did not observe significant burden in any cohort for gene-sets (i)-(iii) (Fig. 2b). However, we still observed a significant variant burden in 22 *FA* genes (OR=2.8; *p*=7.0e-04, 95% CI=1.6–4.7) and 11 *FA* DNA repair genes (OR=2.4; p=0.03; 95% CI=1.1–4.7) in the combined cohort. We next tested how much of this cumulative burden was dependent on all *BRCA2* variants. Even without *BRCA2*, there was still a significant variant burden in *FA* genes (OR=2.6; *p*=5.2e-03, 95% CI=1.4–4.8) in the combined cohort. *FA* core complex does not include *BRCA2*, and remained significant in the combined cohort. FA genes with rare pathogenic variants are listed in Supplementary Table S4.

### *FA* second-hits

We did not observe a case with multiple rare, pathogenic germline *FA* variants. We also did not observe tumor second-hits in any of the cases with a rare, pathogenic germline *FA* variant in the validation cohort (for which we had tumor data), outside of LOH in carriers of *BRCA2* rs11571833 (8/30 individuals) discussed above.

### Effects of smoking

Next, we tested for possible confounding effects of smoking in the discovery cohort (validation cohort controls did not include complete smoking information). Lung SqCC is known to be associated with smoking, and the majority of the individuals in our study were smokers (95.9% in cases and 65.2% in controls). We did not observe noticeable change in the results for any gene-set (Supplementary Fig. S2). Thus, the *FA* genetic effects appear to be independent of smoking history.

### Effects of age and gender

Finally, we tested for confounding effects of gender in all cohorts (gender distribution provided in Supplementary Table S3). We again did not observe a noticeable change in the results for any gene-set (Supplementary Fig. S3). Age, and age plus gender confounding effects in the discovery cohort (validation cohort controls did not include complete age information) also did not reveal a strong effect for any gene-set (Supplementary Fig. S4).

## Discussion

Previous studies have shown that germline variants that increase an individual’s cancer risk occur on a spectrum, from common variants that typically have a modest effect, to rare variants that have high penetrance (23). In this study, we have focused on identifying rare, pathogenic germline variants, which can provide insights into the molecular basis of both familial and sporadic lung SqCC tumorigenesis and a basis for the rational development of personalized prevention strategies for at-risk individuals. For this purpose, we have utilized WES case-control datasets to discover and validate genes with high burden of rare germline variants that confer increased risk for lung SqCC. Ours is the first study that involves joint analyses of rare, disruptive variants that may affect SqCC risk cumulatively as part of a gene or gene-set. While such groupings of single variants based on their cumulative burden on each gene has been shown to increase power to identify new risk genes that drive other diseases (24), they have so far not been used for identifying SqCC risk. Here, we have grouped single variants based on their cumulative burden on each gene, and thus addressed the issue of low power observed in single, recurrent variant studies by a joint analysis of rare, disruptive variants that may affect risk cumulatively as part of a gene. We have further augmented this gain of power by incorporating supporting biological knowledge on functionally related gene groups important in cancer predisposition, another first in SqCC risk studies.

Our study further demonstrates that by using the ClinVar database to restrict analyses of WES data to only those variants known to have a clinical impact, even on other diseases, we can identify important risk genes and gene groups for a disease. Our results support *BRCA2* as a lung cancer risk gene, and we discovered that rare, pathogenic variants in *FA* genes also increase lung SqCC risk.

Within our analyses, we have pursued a resource-conscious approach, leveraging all publicly available germline WES datasets on lung SqCC in TCGA and dbGaP for validation, demonstrating the utility of these databases and especially the importance of analyzing germline DNA from cancer genomics studies. This arguably represents the first successful use of the TCGA germline exome data on sporadic cases to validate a novel set of inherited risk genes for a specific cancer. Indeed, the inherited *BRCA2* and *FA* genetic risk factors were significant in the unselected (sporadic) cohort from TCGA. These observations are in agreement with literature that suggests that genetic predisposition can play a substantial role in the disease onset in sporadic cases, comparable to genetically enriched cohorts (25). For example, recent research has shown that about a quarter of individuals with breast cancer have a risk variant identified from multi-gene sequencing tests (26). In fact, pathogenic variants for which guidelines advise a change in care were detected twice as often in patients who had multi-gene sequencing than in those who only had BRCA1 and BRCA2 analyzed. Our results suggest the development of similar multi-gene genetic testing panel may be useful in helping predict risk of SqCC.

*FA* genes are known to encode for proteins involved in multiple pathways that affect DNA damage repair, particularly inter-strand crosslink, which inhibits DNA replication and transcription and arises from exposure to chemicals found in cigarette smoke. Given that lung SqCC is strongly associated with a history of smoking (e.g. 95% of the cases in this study smoked), it is plausible that in these individuals, if the inter-strand cross-links caused by some of the chemicals in tobacco smoke (e.g. 1,2,3,4-diepoxybutane (DEB) - that are repaired by *FA* genes) are left untreated due to *FA* defects, this may lead to tumorigenesis. These results are consistent with a prior study that has shown that lung adenocarcinomas are associated with common variants in 18 *FA* genes and their combined effects with smoking in a Taiwanese population (27). Pathogenic variants in many of the *FA* genes have also previously been implicated in risk for other cancers. Variants in *BRCA2 (FANCD1)* are known to cause greatly increased risk of breast and ovarian cancer (28,29), as well as increased risk of prostate and pancreatic cancers (19,30). *PALB2 (FANCN)* variants have been associated with pancreatic and breast cancer risk (31–33). In fact, germline variants in several *FA* genes, including *BRIP1 (FANCJ), BRCA1 (FANCS)* and *RAD51C (FANCO)* increase risk for both breast and ovarian cancers (29,34–38) and *FANCM and XRCC2 (FANCU)* variants increase breast cancer risk (39–41). While some of these are commonly tested in clinical genetic settings to evaluate an individual’s risk of cancer, our data suggests the inclusion of all *FA* genes in studies that involve the development risk testing for lung SqCC. Further investigation of the role of this full set of *FA* genes in cancer risk across the spectrum of cancers is also warranted.

## Conclusions

Overall, the findings in this study increase our understanding of SqCC predisposition, and warrant the inclusion of all *FA* genes in current targeted gene sequencing studies offered in the clinic (e.g. MSK-IMPACT) which gather data on many patients that will support the development of a multi-gene genetic diagnostic test that identifies high-risk individuals in the clinic (targeted sequencing can be performed in much larger patient cohorts due to their relatively lower price when compared to exome sequencing). Studies in other cancers have shown that high-risk individuals can benefit from personalized precision medicine based surveillance programs (frequent screening for *BRCA* risk mutation carriers in breast and ovarian cancers), as well as chemoprevention options (daily dose of aspirin for colorectal cancer) in both the affected individuals and their families. While current surveillance options for SqCC with low-dose CT scanning may be risky for individuals with *FA* variants, high-risk individuals will then greatly benefit from the development of intensive surveillance and early detection approaches.

## ACKNOWLEDGEMENTS

This work was supported by a grant to Z.H.G from LUNGevity Foundation and in part through the computational resources and staff expertise provided by Scientific Computing at the Icahn School of Medicine at Mount Sinai. We would also like to thank Dr. Paolo Boffetta for comments on the manuscript.

## AUTHOR CONTRIBUTIONS

M.E., R.J.K. and Z.H.G. wrote the manuscript. Z.H.G supervised the study. R.J.K. performed germline sequence analysis (joint calling). M.E. performed gene burden analysis and other data analyses. R.J.K and Z.H.G. conceived and designed the study.

## COMPETING FINANCIAL INTERESTS

The authors declare no competing financial interests.

